# X-ray irradiation hinders flight ability of *Cydia pomonella* male moths by energy metabolism dysfunction

**DOI:** 10.1101/2025.03.18.644052

**Authors:** Xin-Yue Zhang, Zi-Han Wei, Ping Gao, Yu-Ting Li, Qing-E Ji, Xue-Qing Yang

**Affiliations:** College of Plant Protection, Shenyang Agricultural University, Shenyang 110866, Liaoning, China; Key Laboratory of Economical and Applied Entomology of Liaoning Province, China; Key Laboratory of Major Agricultural Invasion Biological Monitoring and Control, Shenyang 110866, Liaoning, China; Institute of Beneficial Insects, Plant Protection College, Fujian Agriculture and Forestry University, Fuzhou, Fujian, China

**Keywords:** *Cydia pomonella*, Sterile insect technique (SIT), Flight ability, Energy metabolism, ATP

## Abstract

Flight in insects is essential for behaviors such as courtship, foraging, and migration, with energy metabolism serving as its primary energy source. Previous study indicates that X-ray irradiation, even at optimized doses, compromises the flight capacity of male *Cydia pomonella* moths utilized in sterile insect technique (SIT) programs. However, the underlying mechanisms of this phenomenon have not been fully understood. In this study, we report that the disruption of energy metabolism processes contributing to the diminished flight capabilities observed in X-ray irradiated sterile *C. pomonella* moths. Compared to the non-irradiated group, irradiated moths exhibited significantly decreased adenosine triphosphate (ATP) levels. Furthermore, a reduction in the enzymatic activity of key components within carbon metabolism pathways was observed, including citrate synthase (CS) in the tricarboxylic acid cycle (TCA), and glyceraldehyde 3-phosphate dehydrogenase (GAPDH), glycerol 3-phosphate dehydrogenase (GPDH) in glycolysis, and 3-Hydroxyl-CoA dehydrogenase (HOAD) in fatty acid metabolism. Transcriptional analysis revealed downregulation of expression of genes associated with energy metabolism in the primary locomotor tissues post-irradiation, specifically those encoding CS (*CS2*), GAPDH (*Gapdh2*), GPDH (*GPD2*), and HOAD (*HADHA*, *HADH1*, *B0272*, *HADH2*) enzymes. Additionally, exogenous supplementation with decanoic acid and citric acid, the agonist and product of CS respectively, enhanced the TCA cycle, increased energy production, and restored flight capacity in sterile moths. These findings not only confirm that X-ray irradiation disrupts energy metabolism leading to impaired flight, but also offer a potential strategy to overcome this limitation before releasing sterile moths as component of integrated pest management programme using the SIT.

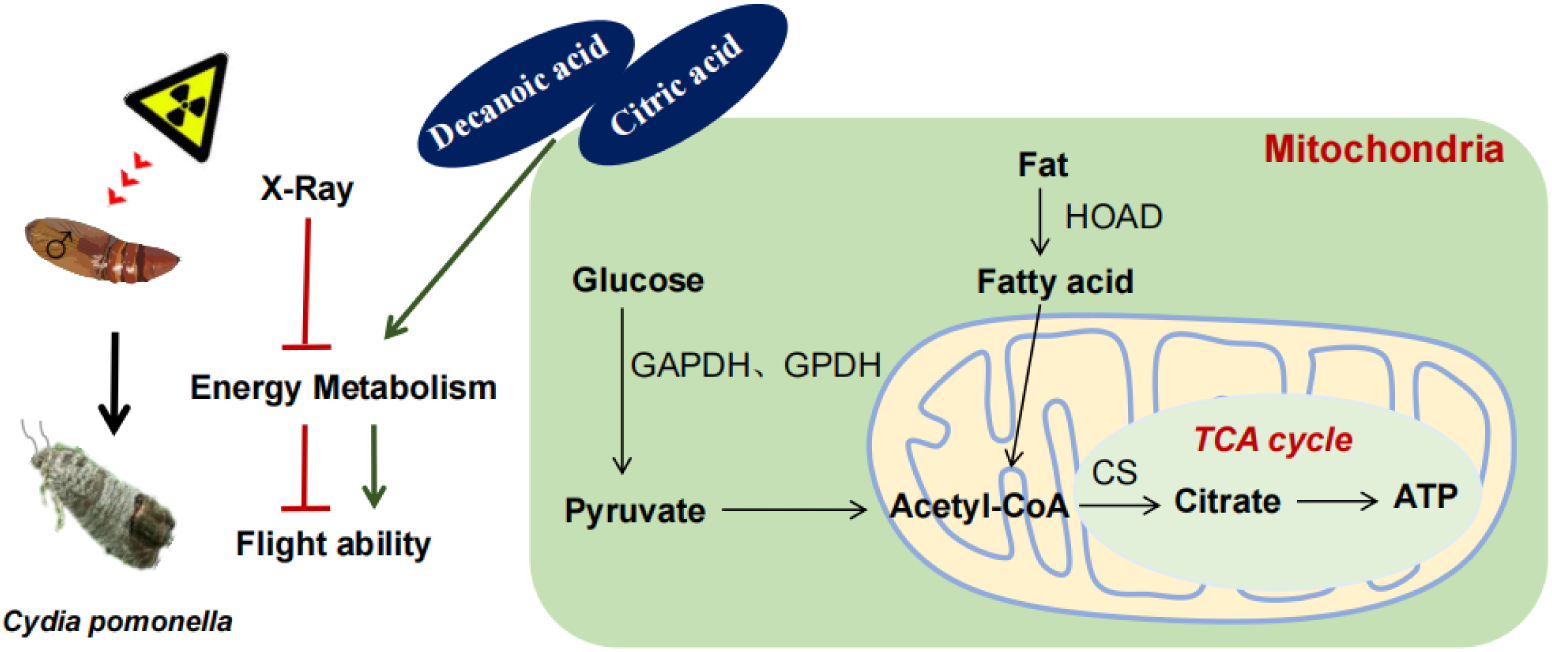
Graphic Abstract.

## Introduction

*Cydia pomonella*, an notorious invasive insect pest that has emerged as a significant threat to fruit production worldwide (Martina et al. 2020). The rapid spread of *C. pomonella* across apple-growing regions on six continents has been facilitated by the exponential growth in global trade and international travel originating from Eurasia since the early 20^th^ century (Ferree et al. 2003). This has resulted in pest establishing its presence in approximately 70 countries, leading to an estimated annual global fruit loss of around US $10 million (Zhang et al. 2024). This pest is considered a major agricultural invasive pest (Yang, 2014; Yang et al. 2022), and has invaded 205 counties within nine provinces in China (http://www.moa.gov.cn/). Current control measures for the *C. pomonella* predominantly rely on chemical interventions (Witzgall et al. 2008), which have raised concerns due to the accumulation of insecticide residues in fruits and emergence of resistance issues (Ju et al. 2021). However, the efficacy of chemical insecticides in managing this pest is hindered by factors such as the overlapping generations and the larvae’ behavior of to tunneling into fruit (Huang et al. 2024; Ju et al. 2021). Thus, there is an pressing necessity to develop new environmentally friendly and sustainable control strategies (Wei et al. 2025).

The Sterile Insect Technique (SIT) is considered a fundamental technology within the area-wide integrated pest management (AW-IPM) (Dyck et al., 2021). By releasing a significant number of sterile males of the target pests, these sterile males engage in competition with wild males, resulting in the production of either no offspring or sterile offspring, leading to a gradual reduction in the pest population (Yamada et al., 2023). A notable milestone in the application of SIT for controlling *C. pomonella* was the successful eradication of populations in the Okanagan Region of Canada through the implementation of the Okanagan-Kootenay Sterile Insect Release (OKSIR) program (Thistlewood and Judd 2019). Nevertheless, despite achieving sterility at the optimal radiation dose, X-ray irradiation exhibits adverse effects on sterile moths particularly in reducing flight performance of *C.pomonella* (Huang et al., 2024b), thereby diminishing the effectiveness of SIT interventions.

The capability of flight is a crucial metric for irradiated sterile male insects upon their release into the natural environment (Diallo et al., 2019). If these sterile insects prove unable to navigate to sheltered areas for survival, procure sustenance, or locate mating sites and female insects through the guidance of sex pheromones, the act of releasing them becomes futile (Dyck et al., 2021). Several studies have shown that exposing sterile insects to radiation can reduce their flight and dispersal capabilities, thereby impeding the effectiveness of releasing them to control pest populations in SIT programs (Smith et al., 2009; Nakamori and Soemori, 1981; Nakamori, 1987; Giancarlo, 2014). Consequently, it is imperative to elucidate the mechanisms responsible for the decreased flight capability induced by X-ray irradiation to develop pertinent enhancement strategies (Huang et al., 2024b). Nonetheless, the specific mechanisms underlying the decline in flight ability in insects due to X-ray radiation remain unknown.

The flight behaviors of insects are impacted by various factors, including physiological characteristics and environmental conditions (Khaliq et al., 2014). These behaviors are additionally governed by a cascade of flight muscle proteins associated with muscle constitution (Yang et al., 2005; Craig and Woodhead, 2006) and enzyme activities linked to energy metabolism (Li et al., 1999; Han et al., 2005; Jenni-Eiermann, 2017). Insect flight is a complex physiological processes, encompassing the conversion of energy substrates, intricate metabolic pathways, and precise hormonal control mechanisms (Li et al., 2023). Energy provisioning is a pivotal determinant influencing the flight capacity of insects, involving carbohydrate and lipid metabolism (Rankin and Burcchsted, 1992), with ATP production serving as the principal energy source. Particularly during the initial flight phase, insects predominantly derive energy from carbon metabolism, wherein the tricarboxylic acid cycle (TCA) serves as a crucial metabolic pathways (Arrese and Soulages, 2010; Van der Horst et al., 1984). Under conditions of sufficient oxygen availability, glycolysis generates compounds that can support the metabolic pathways in most cells (Bar-Even et al., 2012). Pyruvate and nicotinamide adenine dinucleotide (NADH) are transported to the mitochondria, where pyruvate participates in the TCA while NADH is oxidized to NAD^+^ through the electron transport chain, returning to the cytoplasm to aid in glycolysis (Chandel, 2021). Glyceraldehyde 3-phosphate dehydrogenase (GAPDH) plays a pivotal role in glycolysis pathway by converting glyceraldehyde 3-diphosphate (GAP) to glyceraldehyde 1, 3-diphosphate, and producing of NADH using NAD^+^ as a hydrogen receptor (Wang, 2004). The activity of GAPDH is often used as an indicator of glycolysis activity. Glycerol 3-phosphate dehydrogenase (GPDH) is a key enzyme in insects supporting a highly aerobic carbohydrate metabolism state by converting NADH to dihydroxyacetone phosphate (DHAP) and further to glycerol 3-phosphate (G-3-P), while oxidizing NADH to NAD^+^ (Chandel, 2021). This G-3-P shuttle cycle represents as a primary source of energy in the flight muscle of insect, with sugar being the main energy source (Wang 2004). The activity of 3-Hydroxyl-CoA dehydrogenase (HOAD) plays a significant role in fat metabolism levels in insect flight muscles compared to carnitine acyltransferase (Gunn and Gatehouse, 1988). Furthermore, citrate synthase (CS) acts as the initial rate-limiting enzyme in the TCA, catalyzing acetyl-CoA and oxaloacetic acid to produce citric acid and coenzyme A (Verschueren et al., 2019). Citric acid within the cycle regulates energy production by affecting key enzymes in glycolysis, the TCA cycle, gluconeogenesis, and fatty acid synthesis (Iacobazzi and Infantino, 2014). In addition, the TCA is a primary pathway for the complete oxidation and energy provision, and also serve as a crucial biochemical process for the intercoversion of energy substrates, with the activity level within the cycle playing a key role in regulating its function (Wang, 2004). However, the impact of X-ray radiation on insect energy metabolism and its consequent effects on flight ability, as well as the specific key enzymes (genes) involved in this metabolic process, remain to be elucidated.

Hence, the present study aims to determine whether the decrease in flight performance observed in irradiated sterile *C. pomonella* moths is attributed to disturbances in energy metabolism. Furthermore, we seeks to develop an efficient approach for restoring flight ability by addressing the aforementioned metabolic disruptions.

## Materials and methods

### Insects

The *C. pomonella* strain was originated from the Agricultural invasive Biological Control Laboratory of the Institute of Plant Protection, Chinese Academy of Agricultural Sciences. This strain was reared in an incubator (MLR-352H-PC, Panasoni, Matsushita Health Medical Device Co., LTD., Higashi City, Japan) maintained at a temperature of 26 ± 1□, relative humidity of 60% ± 5%, with a light exposure of 16 h followed by 8 h of darkness.

### Irradiation

The X-ray irradiator (SR 100, Changzhou Cerui Instrument Technology Co., LTD., Changzhou City, Jiangsu Province, China) employed in this study was equipped with an a Radcal Accu-Dose+ digitizer featuring a 10 × 6–0.6 CT ion chamber for the purpose of measuring the average dose rate. Male pupae were subjected to 200 Gy of X-ray irradiation the day prior to their emergence and then placed in a cylindrical transparent container (diameter: 2 cm; Length: 6 cm) positioned within the irradiation chamber (size: length: 21 cm; Width: 24 cm; Height: 16 – 40 cm) at a dose rate of 4.3 Gy/ min according to Huang et al (2024b).

### Identification of genes related to energy metabolism in insects

A literature search was conducted to identify genes associated with flight energy metabolism enzymes. Amino acid sequences of Gapdh2, CS, GPD2 and HOAD from *Bombyx mori*, *Plutella xylostella*, *Spodoptera litura*, and other Lepidoptera were obtained from the NCBI database (Table S1) and subsequently employed as query sequences against the *C. pomonella* genome and transcriptome databases. A local blastn query with screening parameter E-value ≤ 10^-5^ was carried out in the *C. pomonella* moth genome database to identify the ORF sequences of related genes. Subsequently, the functional domain corresponding to the gene was searched in the NCBI database for validation.

### Total RNA extraction and cDNA synthesis

RNA was extracted from 10 male adults of fledgling *C. pomonella* moth, which were subjected to either 200 Gy X-ray irradiation or no irradiation, using the TaKaRa MiniBEST Universal RNA Extraction Kit (TaKaRa, Dalian, China). The quantification of RNA samples was conducted with a NanoDrop2000 spectrophotometer (Thermo Fisher Scientific, Waltham, MA, USA). Subsequently, 1 μg of RNA from each treatment was utilized with the PrimeScript RT Reagent Kit (TaKaRa, Dalian, China) to synthesize complementary DNA (cDNA). The resultant cDNA was stored at –20□ for further analysis.

### Real-time quantitative PCR (RT-qPCR) analysis of gene expression

Primers were designed using Primer Premier 5 software (Premier Biosoft International, Palo Alto, CA, USA). The primers used for the RT-qPCR are shown in Table S2. RT-qPCR was performed on a Bio-Rad CFX96 system (Bio-Rad, Hercules, CA, USA) using a 20 μL reaction mixture comprising 1 μL cDNA template, 10 μL TB Green® *Premix Ex Taq*™ (TaKaRa, Dalian, China), 0.8 μL (10 μmol/L) of each primer, and 7.4 μL of double deionized water (ddH_2_O). The reaction condition involved a pre-denaturation step at 95□ for 30 sec, followed by denaturation at 95□ for 5 sec, annealing at 56-61□ for 30 sec, extension at 72□for 30 sec, totaling 40 cycles. Post-reaction, nonspecific amplification was assessed through dissolution curve analysis. The expression of target genes was normalized using *EF-1*α (MN037793) and *RPL12* (MT116775) as internal reference genes (Wei et al., 2021). The relative expression level of each gene was calculated using 2^-ΔΔCt^ method (Livak et al., 2001).

### Quantification of ATP content

ATP content was quantified using the CheKine™ Micro ATP Content Assay Kit (Iacoin Biotechnology Co., LTD., Wuhan, China) following the manufacturer’s instructions. The fundamental concept behind this procedure involves the enzymatic activity of creatine kinase in catalyzing the conversion of creatine and ATP into creatine phosphate. The quantification of creatine phosphate is achieved through phosphomolybdate colorimetry at 700 nm, a method that reacts with the ATP content present. To begin, the tissue sample must be accurately weighed to 0.1 g and combined with 1 mL of deionized water. Subsequently, homogenization of the tissue in an ice bath is necessary, followed by heating the homogenate at 100□ for 5 mins. After centrifugation at 4□ for 15 mins, the upper layer is separated and quantified. For the assay setup, specific reagents are allocated to different wells: the blank well receives 10 μL of Standard, 10 μL of Reagent II, and 30 μL of deionized water. The standard well is treated with 10 μL of Standard, 20 μL of Reagent I, 10 μL of Reagent II, and 10 μL of Reagent III. In the assay well, 10 μL of sample, 20 μL of Reagent I, 10 μL of Reagent II, and 10 μL of Reagent III are combined. Lastly, the control well contains 10 μL of sample, 10 μL of Reagent II, and 30 μL of deionized water. Following thorough mixing, an incubation at 37□ for 30 mins is essential. Subsequently, 200 μL of Working Reagent is added, and the contents are further incubated at 37□ for 20 mins. The absorbance is measured at 700 nm, and the calculations for ATP content are determined by the formulas provided: ATP content (μmol/g fresh weight) = [C_Standard_ × ΔA determination ÷ ΔA standard × V1]÷(W × V1 ÷ V2) = 2 × ΔA determination ÷ ΔA standard ÷ W.

### Quantification of trehalose and citric acid content

The trehalose content was determined using the anthrone method (Zhang et al., 2025) using the Trehalose Content Detection Kit (Beijing Solebao Technology Co., LTD., Beijing, China) according to the manufacturer’s instructions. The quantification of trehalose was conducted through anthrone colorimetry. The procedure involves initially weighing the sample at around 0.1 g and grinding it at room temperature. Subsequently, 1 mL of extract solution is to be added to the sample, which is then to be left at room temperature for 45 mins. Following this, the mixture should be agitated 3 to 5 times before undergoing centrifugation at room temperature to collect the upper layer. The standard product needs to be dissolved in 1 mL of double-distilled water at a concentration of 10 mg/mL, which is then diluted to concentrations of 0.1, 0.08, 0.06, 0.04, 0.02, and 0 mg/mL. To prepare reagent 1, 16 mL of distilled water should be added to each bottle, followed by the slow addition of 64 mL of concentrated sulfuric acid while continuously agitating the contents. After complete dissolution, the reagent is ready for use. For the testing process, 0.25 mL of the standard liquid and 1 mL of the working liquid from an Eppendorf tube are required to be heated in 95□ water for 10 mins, with the tubes tightly covered to prevent water loss, and then allowed to cool naturally to room temperature. Subsequently, 1 mL of colorimetric plate is added, and the absorption value A at 620 nm is measured. The establishment of a standard curve involves plotting the concentration (y) against the corresponding value of the standard sample (x). For the sample analysis, 0.25 mL of the sample is placed in a 1-mL working liquid EP tube, heated in 95□ water for 10 mins with tight covering, cooled to room temperature, and then mixed with 1 mL of colorimetric plate, with the absorption value A at 620 nm measured. The trehalose content y (mg/mL) in the sample can be calculated based on the standard curve, and the trehalose content (mg/g fresh weight) is determined by the formula: trehalose content (mg/g fresh weight) = y ÷ W, where V1 is the volume of the sample, y is the trehalose content in mg/mL, W is the weight of the sample, and V2 is the volume of the working liquid.

Citric acid was extracted from the insect chest for evaluation using a Citric Acid Content Test Kit (Beijing Soleibao Technology Co., LTD., Beijing, China). The reduction of Cr^6+^ to Cr^3+^ by citric acid was monitored at 545 nm, and the citric acid concentration in the sample was subsequently calculated. The tissue specimen needs to be precisely weighed, and approximately 0.1 g of the tissue should be combined with 1 mL of reagent 1 to generate an ice bath homogenate. Subsequently, the homogenate should be centrifuged at 10,000 g at 4□ for a duration of 10 mins, followed by the retrieval of the upper layer for placement on ice for subsequent analysis. The essential reagents for this procedure include distilled water (20 μL), reagent 1 (140 μL), reagent 4 (20 μL), and reagent 5 (20 μL), which are to be introduced into a labeled blank tube. Furthermore, in a measuring tube, 20 μL of the sample, 140 μL of reagent 1, 20 μL of reagent 4, and 20 μL of reagent 5 should be combined. The standard tube should contain 20 μL of the standard, 140 μL of reagent 1, 20 μL of reagent 4, and 20 μL of reagent 5. Upon completion of the mixing step, allow the sample to incubate at room temperature for 30 mins. Subsequently, measure the level of light absorption at a wavelength of 545 nm. The tubes employed in this assay should be properly labeled as the blank tube, determination tube, and standard tube. Calculate ΔA determination by subtracting the absorbance of the determination tube from that of the blank tube, and ΔA standard by subtracting the absorbance of the standard tube from that of the blank tube. Finally, determine the citric acid content quality (μmol/g) using the formula: C standard present × ΔA determination ÷ ΔA standard × V always ÷ W = 2 × ΔA determination ÷ ΔA standard ÷ W.

### Determination of the activity of energy metabolism-related enzymes

The sample was weighed and subjected to a 1:9 (weight: volume) ratio of PBS (PH 7.4) for buffer preparation at a concentration of 0.01 mol/L. Then the samples were homogenized at 4□, followed by centrifugation at 12000 g for 10 mins to collect the supernatant. The original disparity in standards should be lessened by a factor ranging between two to five concentrations. The method comprises the establishment of a control well, a standard well, and a sample well for analysis purposes. The standard substance needs to be dispensed onto the enzyme-coated plate accurately in a 50 μL volume. Subsequently, 40 μL of sample diluent should be introduced into the designated sample well, followed by 10 μL of the sample itself (resulting in a fivefold dilution of the sample). Then carefully dispense the sample into the the base of the enzyme-coated plate well, ensuring that the well walls are not contacted, and gently agitate the contents. The plate is then covered with a sealing film and placed in an incubator at 37□ for 30 mins. The washing solution, concentrated 30 times, should be diluted with 30 parts distilled water for future use. After carefully removing the sealing film, discarding the liquid, and drying the plate, it is ready for further analysis. The procedure involves adding washing liquid to each well of the plate, allowing it to sit for 30 sec, discarding the liquid, and repeating this process five times before drying the plate. Subsequently, 50 μL of enzyme-labelled reagent is added to each well except the blank one, followed by a 30-minute incubation at 37□. The plate is then washed as previously described, followed by the addition of 50 μL of color developer A and 50 μL of color developer B to each well. After gently shaking and mixing, the color developers are developed at 37□ for 10 mins. Termination solution (50 μL) is then added to each well, causing the blue color to change to yellow. The OD of each well is measured at 450 nm, using the blank well as a reference point, within 15 mins of adding the termination solution. A standard curve is then plotted on coordinate paper, with the concentration of the standard substance on the horizontal axis and the OD value of the sample on the vertical axis. The sample’s concentration is determined based on its OD value, multiplied by the dilution factor to certain the actual concentration.

The activity of CS was determined using an ELISA kit (Shanghai Enzyme-Linked Biotechnology Co., LTD., Shanghai, China), utilizing the double antibody sandwich method according to the manufacturer’s instructions. The evaluation of CS levels involved coating purified insect CS antibodies onto microporous plates to create solid phase antibodies. Subsequently, HRP labeled antibodies were added to form an antibody-antigen-enzyme-labeled antibody complex, and the enzymatic activity was assessed through color development using the substrate 3,3’,5,5’-Tetramethylbenzidine (TMB) at an absorbance of 450 nm.

GAPDH activity was determined using the GAPDH enzyme-linked immunoassay kit (Shanghai Enzyme-Linked Biotechnology Co., LTD., Shanghai, China). According to the manufacturer’s instructions, the double antibody sandwich method was employed to quantify GAPDH levels by measuring the characteristic absorption value at 450 nm.

Similarly, glycerin-3-phosphate dehydrogenase (GPDH) activity and 3-hydroxyl-CoA dehydrogenase (HOAD) activity were evaluated using corresponding enzyme-linked immunoassay kits (Shanghai Enzyme-Linked Biotechnology Co., LTD., Shanghai, China) by monitoring the characteristic absorption value at 450 nm.

Lactate dehydrogenase (LDH) enzyme-linked immunoassay kit (Shanghai Enzyme-Linked Biotechnology Co., LTD., Shanghai, China) was used to assess the enzyme activity of LDH according to the manufacturer’s instructions. By measuring the characteristic absorption value at 450 nm using double antibody sandwich method, the LDH activity was evaluated.

### Supplementation with citric acid (CA) and decanoic acid (DA)

Citric acid serves as the catalytic product of CS, while decanoic acid act as the activator of CS (Hara et al., 2021). To evaluate the impact of CS on flight performance post X-ray irradiation, *C. pomonella* moth were externally supplemented with either citric acid (Beijing Solebao Technology Co., LTD., Beijing, China) or decanoic acid (Beijing Solebao Technology Co., LTD., Beijing, China). The experimental protocol involved feeding male *C.pomonella* moth a sucrose solution containing 4 mM citric acid for 48 h within 24 h after emergence.

The decanoic acid supplement, prepared by dissolving 100 mM decanoic acid in DMSO, was administered within the same timeframe. The moths were then randomly divided into 3 groups within 24h: a control group (0 Gy + DMSO), an X-ray irradiation group (200 Gy + DMSO), and a decanoic acid supplement group (200 Gy + decanoic acid). The control moths were fed with a sucrose solution supplemented with DMSO, while the decanoic acid concentration in the sucrose solution for the 200 Gy X-ray irradiation group was adjusted to 250 μM by adding the 100 mM decanoic acid supplement in sucrose solution. The moths were fed the nutrient solution for a duration of 48 h.

### Flight capability analysis

The flight capability was measured using the FXMD-24-USB insect flight information system (Henan Hebi Jiaduoke Industry and Trade Co., Ltd, Hebi, Henan, China) following the protocol outlined in a previous publication (Huang et al., 2024a). Healthy male moths with intact wings that were 3 days post-emergence were positioned on a flat surface using delicate tweezers for scale removal from the upper abdomen with a small brush. A copper wire was fashioned into a circular shape and bend at a 90°angle, then coated lightly with 502 glue. Affix the outer side of the copper wire ring to the upper abdomen of the moth, and apply gentle air pressure to expedite glue solidification. Subsequently, bend the copper wire approximately 1 cm away from the ring to orient the insect’s body perpendicular to the ground. Secure the copper wire onto the flying mill boom, adjusting its position to align the insect’s flight trajectory with the boom. Once positioned, insert the tethered moth’s suspension arm into the flying mill’s base sequentially and ensure its ability to fly post arousal. Activate the flight information system to log pertinent details such as species, age, gender of the test moths, and establish the flight mill’s operation schedule from 21:00 pm to 8:00 am. Upon completion, the system will automatically halt operations, while the data acquisition system records and computes flight duration, distance, speed, and other flight parameters. Additionally, maintain complete darkness in the incubator housing the flying mill, with a temperature of 25 ± 1□ and the relative humidity of 70 ± 5%.

## Data analysis

The statistical significance of gene expression and flight characteristics, including flight duration, distance, and speed of male *C. pomonella* moths, was meticulously analyzed by GraphPad Prism5 (GraphPad Software, SanDiego,CA, USA), *t*-Test to determine if the difference observed in each treatment were statistically significant (\**P* < 0.05; ** *P* < 0.01; \*\*\**P* < 0.001). The results are represented graphically using GraphPad Prism5 software, with all data values expressed as mean ± standard error.

## Results

### X-ray irradiation decreases the ATP content in sterile male *C. pomonella* moths

To evaluate whether the sterile male *C. pomonella* moths following X-ray irradiation impact on the energy metabolism (Figure 1a), we determined the ATP content in 200 Gy of X-ray irradiated and non-irradiated groups. In comparison to non-irradiated control, ATP content in the 200 Gy of X-ray irradiated sterile male moths was significantly decreased (*P*=0.0029, Figure 1 b). This finding suggests that X-ray irradiation decrease ATP production in male *C. pomonella* moths.

**Figure 1.**
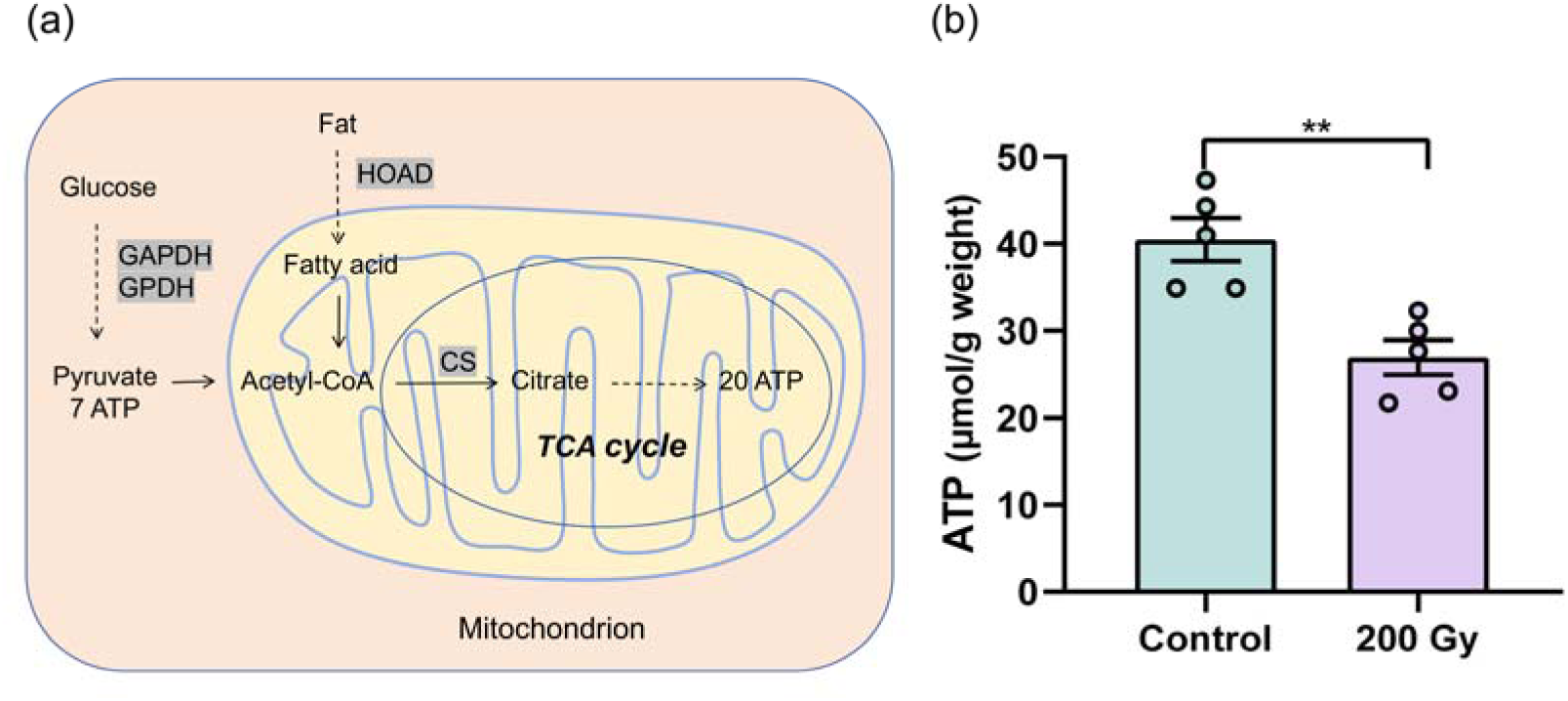
Effects of 200 Gy X-ray irradiation on ATP production in *C. pomonella* moth. (a) Energy metabolism pathway. (b) ATP content in thorax of irradiated and non-irradiated *C. pomonella* males. CS, citrate synthase; GAPDH, glyceraldehyde 3-phosphate dehydrogenase; GPDH, glycerol 3-phosphate dehydrogenase; HOAD, 3-Hydroxyl-CoA dehydrogenase. The data were mean ±SD. The asterisk indicates significant differences as determined by student’s *t*-test (\**P*<0.05, \*\**P*<0.01, \*\*\**P*<0.001). n.s. indicates no significant difference.

### X-ray irradiation inhibits key enzymes involved in energy metabolism

To establish a correlation between the reduction in ATP content induced by X-ray irradiation-induced and potential dysfunction in energy metabolism, we evaluated the performance of critical enzymes participating in this metabolic pathway. The standard curve of concentration and OD value of GAPDH, GPDH, HOAD, LDH, CS are shown in Figure S1. Results revealed a substantial decrease in CS activity in the whole body (Figure 2a), thorax (Figure 2b), and leg (Figure 2c) of sterile male moths following X-ray exposure at 200 Gy, compared with the control group. Genes involved in energy metabolism in *C. pomonella* were identified by homology comparison (Table S3). Similarly, the activities of GAPDH (Figure 2 d-f), GPDH (Figure 2 g-i), and HOAD (Figure 2 j-l) in the whole body, thorax, and leg were notably reduced post-irradiation, unlike LDH activity, which showed no variance between the irradiated and non-irradiated groups (Figure S2). These findings suggest that exposure to 200 Gy X-ray irradiation hampers energy metabolic processes and interferes with the function of associated enzymes.

**Figure 2.**
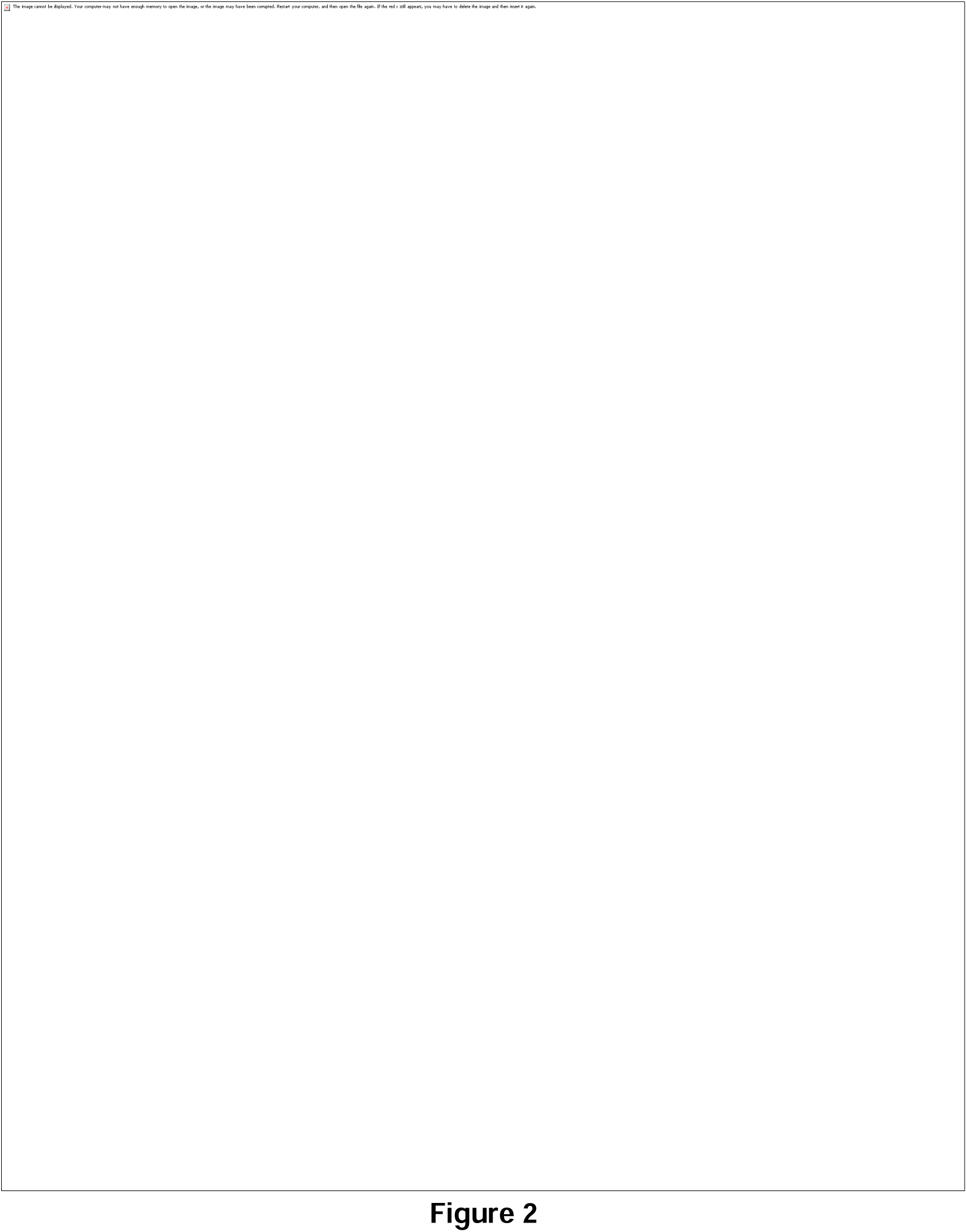
Effects of 200 Gy X-ray irradiation on enzymes related to energy metabolism in *C. pomonella* moth. Activity of citrate synthase (CS) in the whole body (a), thorax (b), and leg (c) of male *C. pomonella* moths irradiated with or without 200 Gy X-ray. Activity of GAPDH in the whole body (d), thorax (e) and leg (f) of *C. pomonella* moths irradiated with or without 200 Gy X-ray. Activity of GPDH in the whole body (g), thorax (h) and leg (i) of *C. pomonella* moths irradiated with or without 200 Gy X-ray. Activity of HOAD in the whole body (j), thorax (k) and leg (l) of *C. pomonella* moths irradiated with or without 200 Gy X-ray. The data were mean ±SD. The asterisk indicates significant differences as determined by student’s *t*-test (\**P*<0.05, \*\**P*<0.01, \*\*\**P*<0.001). n.s. indicates no significant difference.

### X-ray irradiation alters carbon metabolic intermediates

Furthermore, citric acid levels displayed a significant decrease (Figure 3a), while trehalose content exhibited a notable increase (Figure 3b) in the X-ray irradiated group compared to the non-irradiated control. These findings suggest that 200 Gy X-ray irradiation alters carbon metabolic intermediates. The increase in trehalose levels could potentially be attributed to compromised carbon metabolism induced by X-ray exposure, leading to a decrease in its utilization.

**Figure 3.**
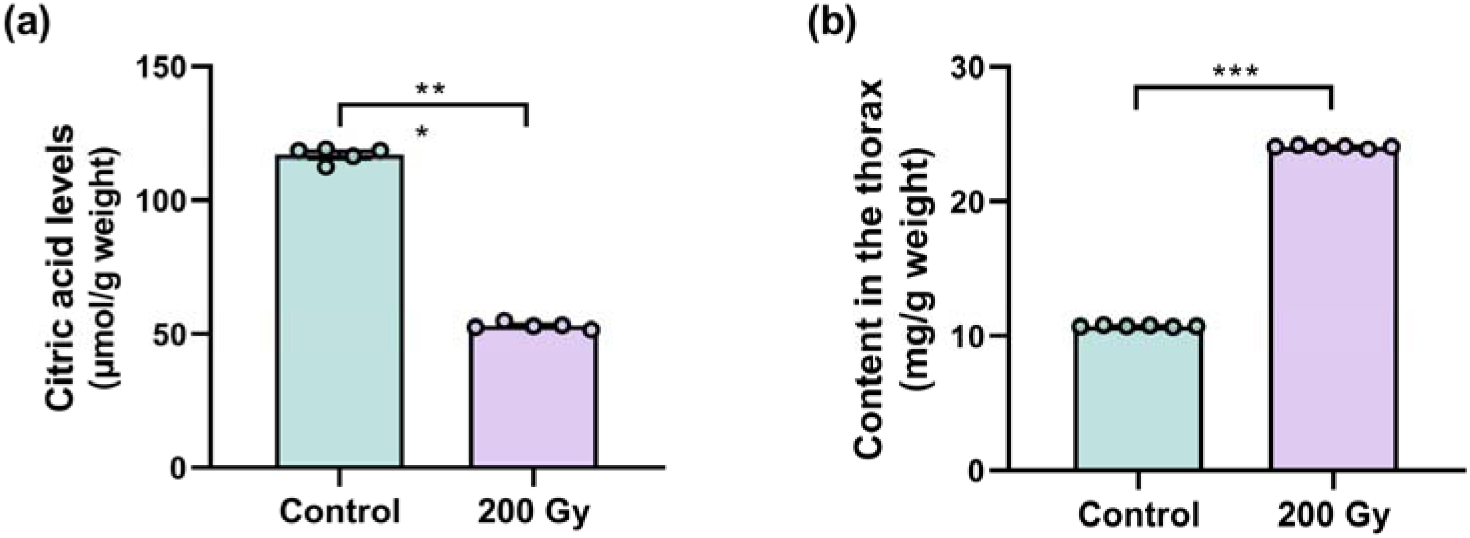
200 Gy X-ray irradiation affected carbohydrate metabolism of *Cydia pomonella* moth. (a) Citric acid content in the thorax of male *C. pomonella* moths. (b) Trehalose content in the thorax of male *C. pomonella* moths. The data were mean ±SD. The asterisk indicates significant differences as determined by student’s *t*-test (\**P*<0.05, \*\**P*<0.01, \*\*\**P*<0.001). n.s. indicates no significant difference.

### X-ray irradiation downregulates the expression of key genes associate with energy metabolism

To further elucidate the molecular mechanism of X-ray irradiation on energy metabolism, we identified and analyzed the expression levels of key genes involved in this metabolic pathway by RT-qPCR. A total of 1 gene in CS (*CS2*), GAPDH (*Gapdh2*), and GPDH (*GPD2*) each, and 4 genes in HOAD (*HADHA*, *HADH1*, *B0272*, *HADH2*) were identified in *C. pomonella* (Table S3). Gene expression analysis results indicated that the expression level of *CS2,* involved in the citric acid cycle, the last stage of carbon metabolism, was notably reduced in the whole body (Figure 4a) and thorax (Figure 4b) compared to the control group, while no significant difference was found in the leg (Figure 4c). Furthermore, after exposure to 200 Gy X-ray irradiation, the expression of glycolysis-related gene *Gapdh2* in the whole body significantly decreased (Figure 4d), while expression was not detected in thorax and leg. Additionally, the expression of another glycolysis-related gene *GPD2* showed a significant decrease in expression in the whole body (Figure 4e), with no difference in the thorax (Figure S3), and an increase in the leg (Figure S3). Moreover, the expression levels of fatty acid metabolism-related genes *HADHA*, *HADH1*, *B0272*, *HADH2* in the whole body were significantly reduced (Figure 4f-4i), with increased expression levels in the thorax (Figure S4), while no significant difference or a decrease in expression levels was observed in the leg (Figure S4). These results suggest that 200 Gy X-ray irradiation can significantly impede or disrupt the normal energy metabolism of *C. pomonella* moths.

**Figure 4.**
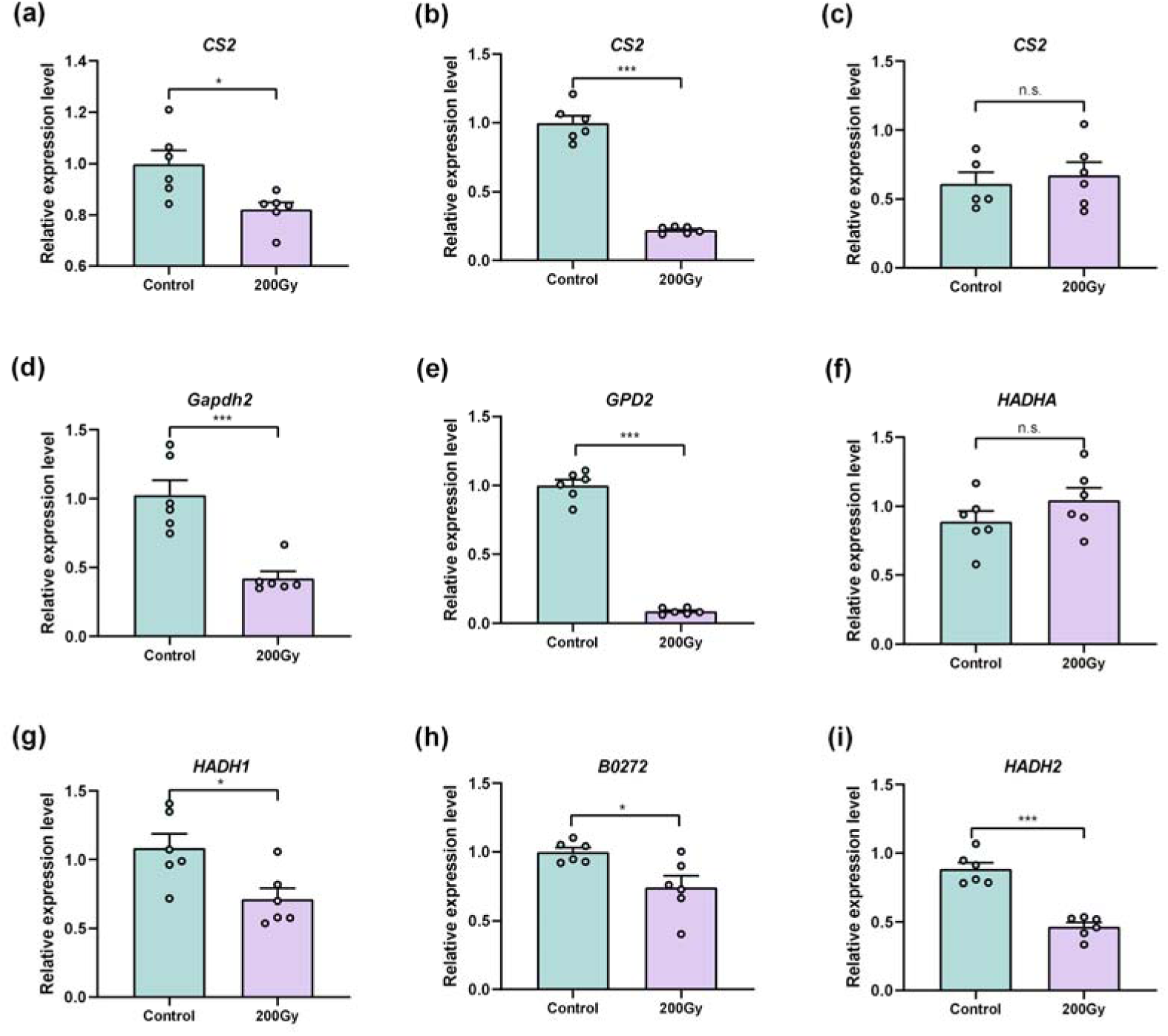
Effect of 200 Gy X-ray irradiation on expression of key genes in energy metabolism in *C. pomonella* moth. The expression level of *CS2* in the whole body (a), thorax (b) and leg (c) of male *C. pomonella* moths irradiated with or without 200 Gy X-ray. The expression level of *Gapdh2* (d), *GPD2* (e), *HADHA* (f), *HADH1* (g), *B0272* (h), *HADH2* (i) in the whole bodyof male *C. pomonella* moths irradiated with or without 200 Gy X-ray. The data were mean ±SD. The asterisk indicates significant differences as determined by student’s *t*-test (\**P*<0.05, \*\**P*<0.01, \*\*\**P*<0.001). n.s. indicates no significant difference.

### Exogenous supplement improves the activity of CS and restores the level of energy metabolism

Supplementation of sterile moths with decanoic acid, an activator of CS, and citric acid, a product of CS, was conducted to investigate the potential for enhancing CS activity in restoring energy metabolism. Results showed that exposure to 200 Gy X-ray irradiation led to a reduction in citric acid content, which was significantly counteracted by supplementation with decanoic acid (Figure 5a). Additionally, supplementation with decanoic acid was found to upregulate the relative expression of *CS2* (Figure 5b) and enhance CS enzyme activity (Figure 5c) in X-ray irradiated males compared to non-irradiated controls. Notably, sterile males with decanoic acid supplementation exhibited significantly higher ATP levels, the final product of carbon metabolism, when compared to the unsupplemented group (Figure 5d). Similarly, supplementation of irradiated males with citric acid also led to increased ATP levels (Figure 5e). Previous study has documented that *myosin heavy chain* is flight-related gene in insect (Huang et al., 2024b). RT-qPCR analysis shows that exogenous supplementation with citric acid and capric acid also significantly increased *myosin heavy chain* expression in irradiated sterile male moths (Figure 5f). These results indicate that supplementation with citric acid or decanoic acid can mitigate the adverse impacts of X-ray irradiation on energy metabolism of *C. pomonella* moths.

**Figure 5.**
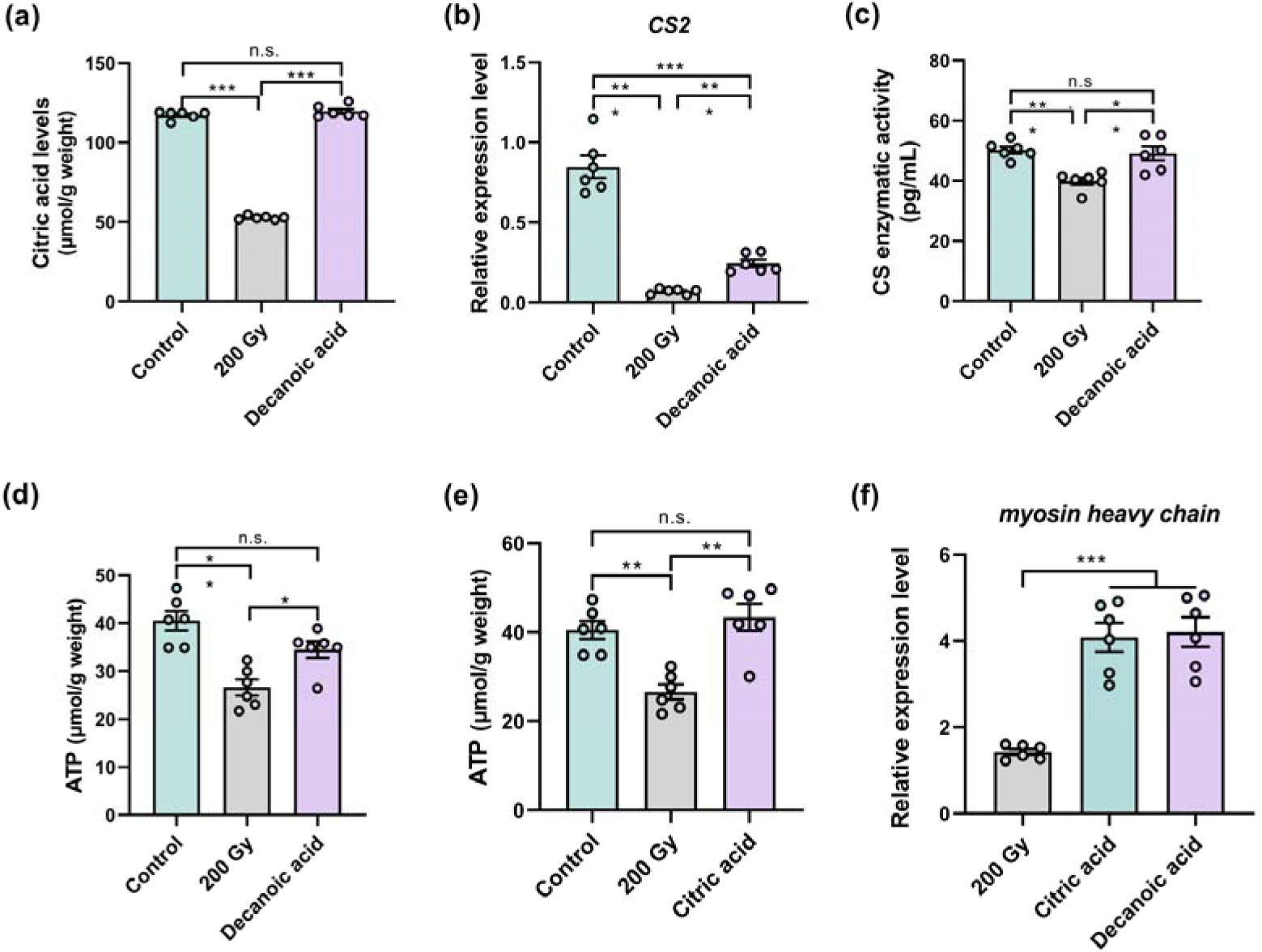
Exogenous supplementation of citric acid or decanoic acid can save energy metabolism after X-ray irradiation in *C. pomonella* moth. (a) Effect of administering exogenous decanoic acid on the citric acid levels in X-ray irradiated *C. pomonella* moths. (b) RT-qPCR analysis of the expression level of *CS2* gene after supplementing decanoic acid in X-ray irradiated *C. pomonella* moths. (c) Effect of administering exogenous decanoic acid on enhances the activity of citrate synthetase in X-ray irradiated *C. pomonella* moths. (d) ATP levels after decanoic acid supplementation in X-ray irradiated *C. pomonella* moths. (e) ATP levels after citric acid supplementation in X-ray irradiated *C. pomonella* moths. (f) RT-qPCR analysis of the expression level of *myosin heavy chain* gene after supplementing decanoic acid in X-ray irradiated *C. pomonella* moths. The asterisk indicates significant differences as determined by student’s *t*-test (\**P*<0.05, \*\**P*<0.01, \*\*\**P*<0.001). n.s. indicates no significant difference.

### Exogenous supplement restores the flight performance of sterile moths

To determine whether restoring the function of citrate synthase would mitigate the adverse effects of X-ray irradiation on flight performance of *C. pomonella* moth, the flight duration, speed, distance were monitored. Results indicated that 200 Gy of X-ray irradiation did not alter the flight speed of sterile male moths, including the max velocity (Figure 6a) and mean velocity (Figure 6b), but it decreased the flight duration (Figure 6c), consequently resulting in a shortened flight distance (Figure 6d). Exogenous application supplementation with citric acid and decanoic acid both significantly prolonged the flight duration (Figure 6c) of irradiated sterile male moths, thereby expanding the flight distance (Figure 6d). These results indicate that external provision of citric acid or decanoic acid can mitigate the adverse impacts of X-ray irradiation on the flight performance of *C. pomonella* moths, probably through restoring their energy reserves.

**Figure 6.**
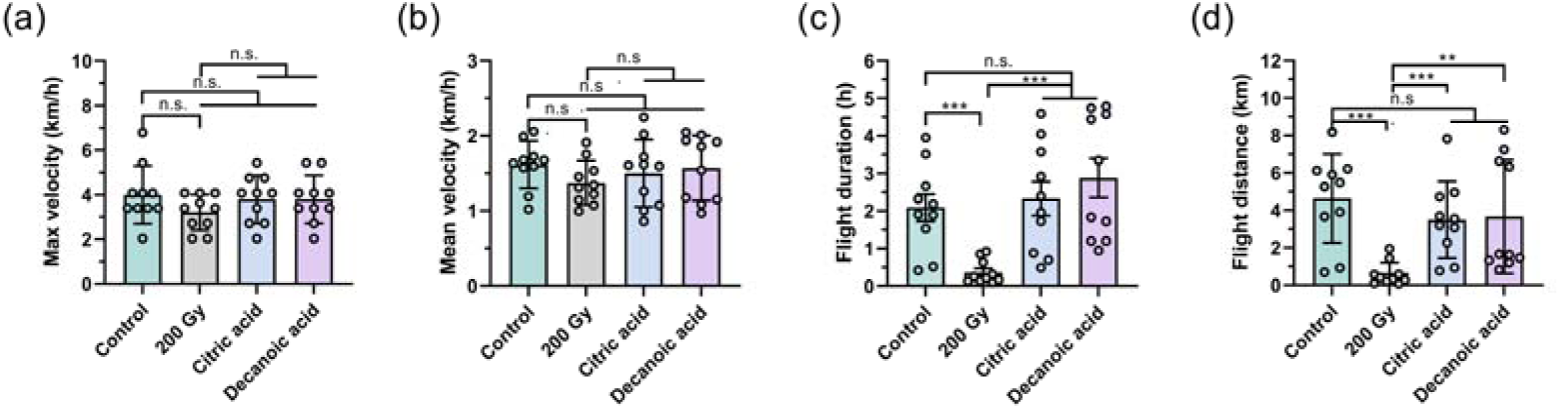
Supplementation with citric acid or decanoic acid reversed the adverse effects of X-ray irradiation on the flight ability of the *C. pomonella* moth. Effects of administering exogenous citric acid or decanoic acid on the maximum (a) and mean (b) flight velocity of X-ray irradiated *C. pomonella* moths. (c) Effects of administering exogenous citric acid or decanoic acid on the flight duration of X-ray irradiated *C. pomonella* moths. (d) Effects of administering exogenous citric acid or decanoic acid on the flight distance of X-ray irradiated *C. pomonella* moths. The data were mean ±SD. The asterisk indicates significant differences as determined by student’s *t*-test (\**P*<0.05, \*\**P*<0.01, \*\*\**P*<0.001). n.s. indicates no significant difference.

## Discussion

The ability of insect to fly is essential for them to locate shelter, find sustenance, and navigate towards the females using sex pheromones (Wagner and Liebherr, 1992; Holland et al., 2006; Dyck et al. 2021). Previous studies have shown that subjecting males *C. pomonella* to an optimal X-ray irradiation dose of 200 Gy significantly impairs their flight capabilities (Huang et al., 2024a), thus greatly reducing the efficacy of using SIT for pest control. In this study, we illustrated that the diminished flight capacity in male *C. pomonella* moths following irradiation is a result of compromised energy metabolism and inadequate energy supply. By externally administering compounds to enhance CS activity, energy metabolism can be restored, consequently improving the flight performance of sterile male *C. pomonella* moths.

A decrease in flight performance of insect may lead to a reduction in encounters with potential mates, resulting in a decline in mating frequency, potentially impacting the efficacy of SIT programs (Wagner and Liebherr, 1992; Holland et al., 2006; Dyck et al., 2021). Several studies have indicated a reduction in flight capabilities among sterile insects following irradiation (Nakamori, 1987; López-Martínez et al., 2014). For instance, one experiment revealed a decline in the flight ability of *Dacus cucurbitae* as the dosage of γ-ray irradiation escalated from 7 to 30 KR (Nakamori, 1987). Additionally, another study observed a decrease in the flight performance of male *Anastrepha suspensa* exposed to γ-ray irradiation (López-Martínez, 2014). In this study, the flight capacity of *C. pomonella* moths was notably impacted by 200 Gy X-ray irradiation, as evidenced by a significant decrease in flight distance and duration compared to non-irradiated counterparts. These findings are consistent with previous research by Huang et al. (2024b) involving 366 Gy X-ray irradiation. The current study conducted statistical analysis from 21:00 p.m. to 08:00 a.m, totaling 11 hours. Male moths exposed to 200 Gy X-ray irradiation exhibited a mean flight speed of 1.5 km/h, flight duration of 2 h, and flight distance of 4 km. The experiment showed a decrease in flight speed by 0.2 km/h, along with reductions of 0.1 h in flight duration and 0.1 km in flight distance post-irradiation. Huang et al. (2024b) reported similar findings with statistical analysis over 8 h, where unirradiated male moths had an average flight speed, duration, and distance of 1.18 ± 0.09 km/h, 53.82 ± 7.17 min, and 1.06 ±0.28 km, respectively. When exposed to 366 Gy X-ray irradiation, male moths displayed an average flight speed of 0.48±0.06 km/h, flight duration of 5.28 ± 7.63 min, and flight distance of 65.41±33.19 m (Huang et al., 2024b).

Discrepancies in flight metrics between this study and Huang et al. (2024b) could potentially be attributed to variations in statistical times and irradiation doses. Interestingly, while the mean flight velocity of the moths remained unaffected by X-ray irradiation in this study, contrary to observations following 366 Gy X-ray irradiation (Huang et al. 2024b), it is plausible that only high dose of X-ray irradiation significantly impact flight speed. Notably, the max flight velocity of male moths subjected to X-ray irradiation was comparable to that of the control group, indicating that X-ray irradiation primarily influences the flight endurance of *C. pomonella* moths.

The experimental results demonstrated that X-ray irradiation suppressed the expression of key genes and enzymes involved in the carbon metabolism pathway. This suppression led to the obstruction of carbohydrate catabolism and exerted deleterious effects on metabolic pathways, including glycolysis and the TCA cycle (Gao et al., 2024). Given the high energy expenditure required during insect flight (Harrison and Roberts, 2000), the reduction in ATP levels in the experimental results may explain the significant reduction in flight persistence. Gao et al. (2024) also reported that the impaired flight ability of insects caused by stress was due to impaired carbon metabolism and insufficient energy supply. Carbohydrates are the primary source of energy for flight in lepidoptera insects, utilizing sugars directly from its food during flight (Rankin and Burcchsted, 1992). Previous findings indicated that there was no significant change in the overall sugar and glycogen levels of sterile males after exposure to irradiation (unpublished dada). However, this study showed that there was a notable increase in trehalose content, possibly resulting from the inhibition of carbohydrate breakdown due to X-ray irradiation. These results contrast with the research by Gao et al. (2024) on bumblebees, where glucose and trehalose levels rose under cadmium stress. In our study, while the total sugar and glycogen levels in *C. pomonella* moths remained steady post-irradiation, trehalose levels increased, potentially influenced by different stress factors. It is important to note that the effects of cadmium stress differ from the potential harm caused by X-ray irradiation in insects. Notably, key metabolic enzymes such as GAPDH, GPDH, HOAD, and CS exhibited inhibited activity, indicating compromised carbon metabolism. Conversely, LDH activity remained unchanged, aligning with previous study by Li et al. (1999), suggesting a minor role of LDH in insects in response to X-ray irradiation stress. X-ray exposure led to decreased citric acid levels in the body, consistent with reduced CS activity, in agreement with the findings of Gao et al.(2024). Notably, glycolysis from 1 glucose molecule can yield 7 ATP, while complete TCA oxidation can produce 20 ATP (Chandel, 2021). Moreover, it has been observed that the inhibition of CS activity reaches its peak level subsequent to exposure to 200 Gy X-ray irradiation. These findings suggest that X-ray irradiation could potentially disrupt carbohydrate metabolism by influencing the functionality of CS, thereby potentially compromising the physiological capabilities related to flight.

CS facilitates the conversion of acetyl-CoA and oxaloacetic acid into citric acid (Verschueren et al., 2019). Within the cell, citric acid functions as a regulator, influencing energy production and modulating various metabolic enzymes involved in glycolysis, the TCA cycle, gluconeogenesis, and fatty acid synthesis (Iacobazzi and Infantino, 2014). Furthermore, decanoic acid was observed to enhance citric acid synthase activity in mitochondria (Hughes et al., 2014). Previous studies by Hara et al (2021) have shown that supplementing citric acid can significantly increase metabolite levels associated with the TCA cycle in rats. Our research similarly reveals that administering exogenous citric acid or decanoic acid post X-ray exposure can partially improve the flight capabilities of *C. pomonella* moths back to a normal level. Flightin, myosin, light chain, and myosin heavy chain are contractile proteins essential for insect flight performance (Craig and Woodhead, 2006). The observed enhancement is likely due to the ability of these compounds to upregulate the expression of flight-associated proteins. Specifically, the myosin heavy chain protein, a key component of the thick filaments in the myogenic fibers of insect flight muscles, is critical for insect flight kinematics (O’Donnell and Bernstein, 1988; Huang et al., 2024b). We observed a significant increase in the expression of the *myosin heavy chain* gene in 200 Gy X-ray-irradiated males following exogenous supplementation with decanoic acid and citric acid, relative to the control group. These results suggest that exogenous supplementation with decanoic acid and citric acid enhances CS activity, elevates citrate levels, accelerates energy metabolism, and increases ATP production, thereby restoring the flight capacity of sterile moths. These outcomes differ from the findings of Gao et al. (2024) involving bumblebees under cadmium stress, potentially owing to varying levels of damage caused by distinct stressors, resulting in differing recovery rates. Jiang et al. (2019) demonstrated that starvation in *Helicoverpa armigera* markedly suppressed respiratory and energy metabolism, accompanied by a significant downregulation of glycolytic enzyme gene expression. These findings align with our observations, suggesting a consistent inhibitory effect of stress on energy metabolism-related pathways. Kuo et al (2023) reported that elevated temperatures compromised the energy metabolism of bumblebees, diminishing energy allocation to flight muscles, thereby reducing wingbeat frequency and flight distance. This result consists with our findings, which indicate that stress impairs energy metabolism, leading to reduced flight distances in insects. Given the paucity of literature on the effects of X-ray irradiation and other environmental stressors on insect energy metabolism, our findings do not yet allow us to determine whether environmental stressors universally induce energy metabolism dysfunction, which could subsequently impact insect fitness traits such as flight. Nevertheless, our research suggests that elucidating the mechanisms underlying the negative effects of environmental stressors on insects could provide a scientific basis for targeted improvement of these traits.

## Conclusion

Collectively, we evaluated the impact of X-ray irradiation on flight performance and energy metabolism of *C. pomonella* moth. Our results support the conclusion that X-ray irradiation hinders the flight capability of the *C. pomonella* moth by suppressing energy metabolism. Furthermore, it was observed that citrate synthase could potentially serve as a biomarker affecting energy metabolism. The restoration of CS function through supplementation with citric acid or decanoic acid was shown to activate the TCA cycle, elevate energy production (including ATP levels), and subsequently recover flight endurance and distance. As a result, this approach counteracts the adverse effects of X-ray exposure on flight capacity. These findings offer novel insights into the adverse repercussions of X-ray irradiation on insect flight and present valuable implications for the implementation of SIT in managing *C. pomonella* populations.

## Authors’ contributions

**Xin-Yue Zhang**: Performed the experiments, data curation, and writing the manuscript; **Zi-Han Wei**: data curation and revised the manuscript; **Ping Gao & Yu-Ting Li**: Data curation and revised the manuscript; **Qing-E Ji**: Investigation, supervision, and revised the manuscript; **Xue-Qing Yang**: Design the experiments, validation, Writing, revised the manuscript, and funding acquisition.

## Acknowledgements

This work was supported by the National Key R&D Program of China (2021YFD1400200) and Liaoning Province Science Fund for Distinguished Young Scholars (2024JH3/50100027).

