## Supplementary Table 1-Table3, Figure 1-Figure 4 for "X-ray irradiation hinders flight ability of *Cydia pomonella* male moths by energy metabolism dysfunction"

**Table S1. Genes associated with energy metabolism in Lepidoptera were derived from the NCBI database.**

| **Enzymes** | **Gene Bank** | **Organism** | **Description** |
| --- | --- | --- | --- |
| **HOAD** | | | |
|  | XP_061719020 | *Cydia pomonella* | hydroxyacyl-coenzyme A dehydrogenase, mitochondrial-like |
|  | ADW77640 | *Bombyx mori* | 3-hydroxyacyl-CoA dehydrogenase |
|  | XP_028041500 | *Bombyx mandarina* | hydroxyacyl-coenzyme A dehydrogenase, mitochondrial-like |
|  | XP_022822785 | *Spodoptera litura* | hydroxyacyl-coenzyme A dehydrogenase, mitochondrial-like |
|  | KAG7312510 | *Plutella xylostella* | hypothetical protein JYU34_002031 |
|  | KAF9421241 | *Spodoptera exigua* | hypothetical protein HW555_002713 |
|  | XP_035444302 | *Spodoptera frugiperda* | hydroxyacyl-coenzyme A dehydrogenase, mitochondrial |
| ***GAPDH*** | | | |
|  | NP_001037386 | *Bombyx mori* | glyceraldehyde-3-phosphate dehydrogenase |
|  | CAD33827 | *Plutella xylostella* | glyceraldehyde-3-phosphate dehydrogenase |
|  | XP_022822680 | *Spodoptera litura* | glyceraldehyde-3-phosphate dehydrogenase |
|  | XP_035443774 | *Spodoptera frugiperda* | glyceraldehyde-3-phosphate dehydrogenase |
|  | KAF9424578 | *Spodoptera exigua* | hypothetical protein HW555_000389 |
|  | QPZ44475 | *Lymantria dispar* | glyceraldehyde-3-phosphate dehydrogenase |
| ***GPDH*** | | | |
|  | ACV95335 | *Bombyx mori* | glycerol-3-phosphate dehydrogenase |
|  | XP_028034134 | *Bombyx mandarina* | glycerol-3-phosphate dehydrogenase, mitochondrial |
|  | XP_061707611 | *Cydia pomonella* | glycerol-3-phosphate dehydrogenase, mitochondrial isoform X1 |
|  | XP_022820672 | *Spodoptera litura* | glycerol-3-phosphate dehydrogenase, mitochondrial isoform X1 |
|  | XP_026726806 | *Trichoplusia ni* | glycerol-3-phosphate dehydrogenase, mitochondrial isoform X1 |
| ***CS*** | | | |
|  | XM_061874062 | *Cydia pomonella* | probable citrate synthase 2, mitochondrial |
|  | XM_063537095 | *Cydia fagiglandana* | probable citrate synthase 2, mitochondrial |
|  | XM_063981477 | *Ostrinia nubilalis* | probable citrate synthase 2, mitochondrial |
|  | XM_063776798 | *Cydia splendana* | probable citrate synthase 2, mitochondrial |

**Table S2. The primers used for the reverse-transcription quantitative polymerase chain reaction (RT-qPCR).**

| Primer name | Sequence (5'-3') |
| --- | --- |
| HADH1-qPCR-F | TGACGCCACAGCCGAAGA |
| HADH1-qPCR-R | CACCAGCCGGTCCAACACT |
| HADH2-qPCR-F | GTGCCGTATATCGCTGAGG |
| HADH2-qPCR-R | AGGATGAACTTGTTGGTGTCG |
| B0272-qPCR-F | TCGGCGGTCTCCACTTCTTC |
| B0272-qPCR-R | GCTTTGCCCCACTCCATCAT |
| HADHA-qPCR-F | GCGGGCTAGAGACTGCTCTG |
| HADHA-qPCR-R | TCGCTTTATCTGCCTTTACGG |
| Gapdh2-qPCR-F | GTTCACCAGCGTGGAAAAGG |
| Gapdh2-qPCR-R | GCGCCAGGCAGTAAAGGG |
| GPD2-qPCR-F | CGGTAAGGAATGGGATGTAAAAGC |
| GPD2-qPCR-R | TCAGCCAGGGCAGGAAGAAA |
| CS2-qPCR-F | CCACTCTGGCGTCCTTCTACA |
| CS2-qPCR-R | CGCGACCACACCATCTGC |
| myosin heavy chain-qPCR-F | CCGTAACGACAACTCTTCCC |
| myosin heavy chain-qPCR-R | CGACAAAATGCACTTTCCCT |

**Table S3. Genes involved in energy metabolism identified by homology comparison.**

| **Genes** | **Genbank number** | **Organism** | **Description** |
| --- | --- | --- | --- |
| *HADH1* | XM_061863038 | *Cydia pomonella* | Hydroxyacyl-coenzyme A dehydrogenase, mitochondria |
| *HADH2* | XM_061863036 | *Cydia pomonella* | Hydroxyacyl-coenzyme A dehydrogenase, mitochondrial |
| *B0272* | XM_061863037 | *Cydia pomonella* | Probable 3-hydroxyacyl-CoA dehydrogenase B0272.3 |
| *HADHA* | XM_061851275 | *Cydia pomonella* | Trifunctional enzyme subunit alpha, mitochondrial |
| *Gapdh2* | XM_061863454 | *Cydia pomonella* | Glyceraldehyde-3-phosphate dehydrogenase 2 |
| *GPD2* | XM_061851629 | *Cydia pomonella* | Glycerol-3-phosphate dehydrogenase, mitochondrial |
| *CS2* | XM_061874062 | *Cydia pomonella* | Probable citrate synthase 1, mitochondrial |

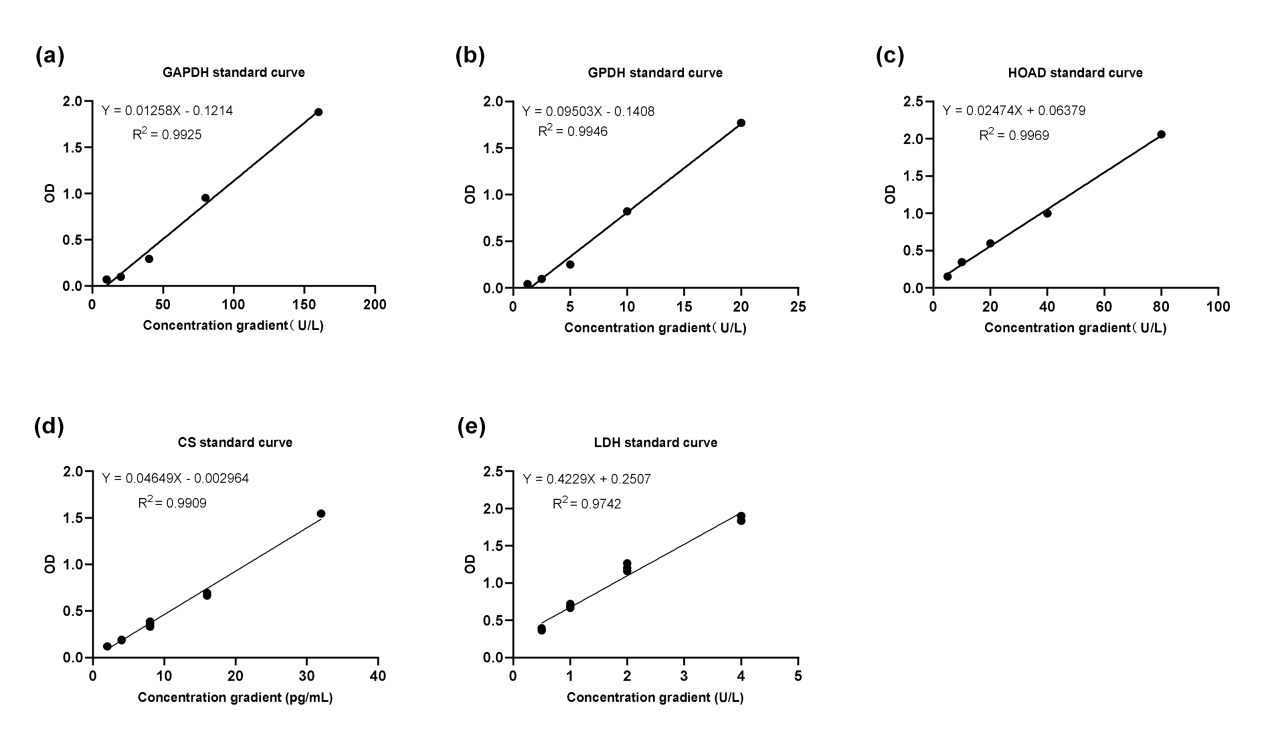

**Figure S1. Standard curve of concentration and OD value of energy metabolism-related enzymes in *C. pomonella*.** (a) Standard curve of *GAPDH*; (b) Standard curve of *GPDH*; (c) Standard curve of *HOAD*; (d) Standard curve of *CS*; (e) Standard curve of *LDH.*

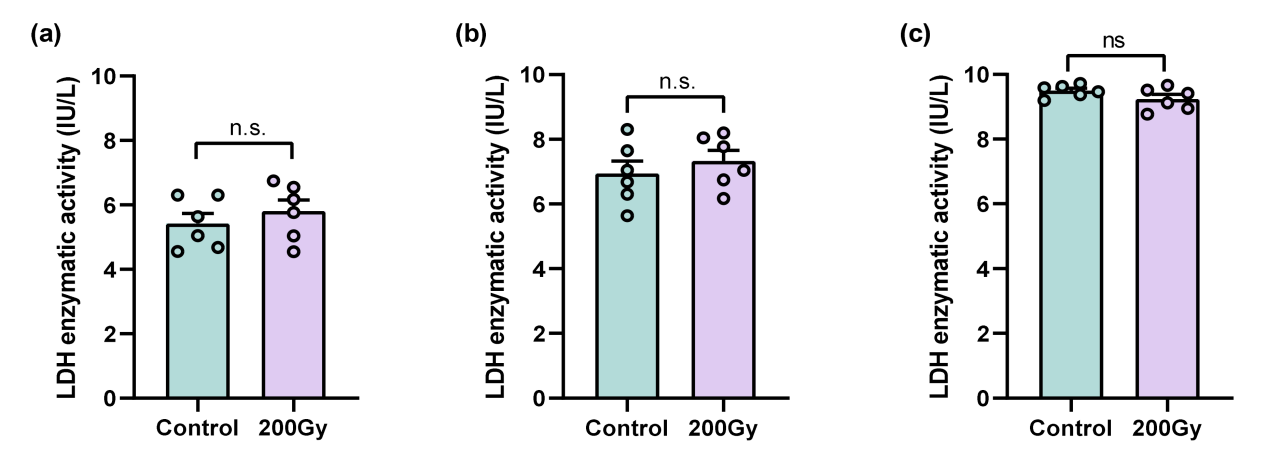

**Figure S2. Effect of 200 Gy X-ray irradiation on the activity of lactate dehydrogenase (LDH) in *C. pomonella* moth.** (a) LDH activity in the whole body (a), thorax (b), and leg (c) of male *C. pomonella* moths irradiated with or without 200 Gy X-ray. The data were mean ±SD. n.s. indicates no significant difference determined by student’s *t*-test (*P*<0.05).

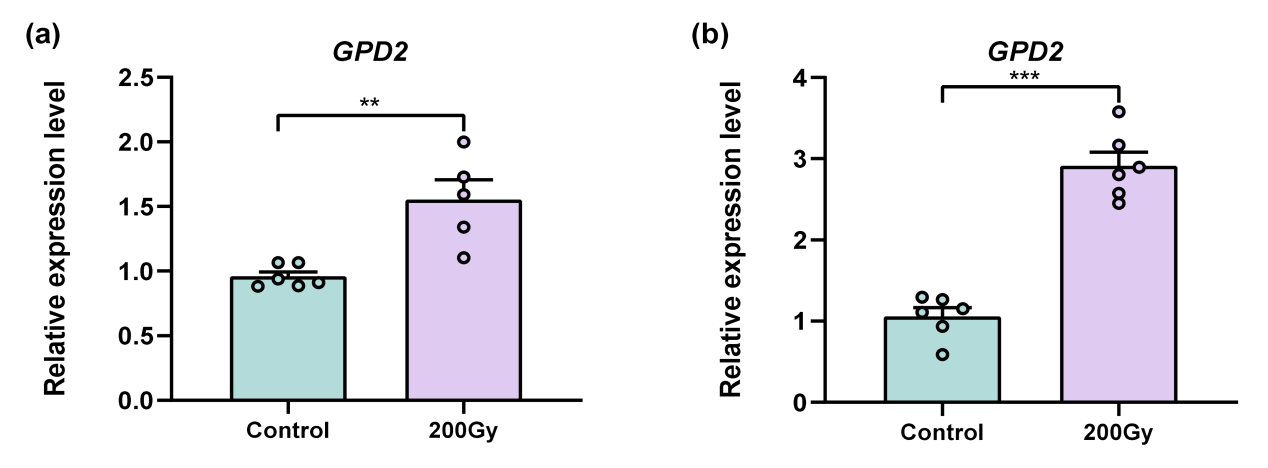

**Figure S3. Effect of 200 Gy X-ray irradiation on expression of *GPD2* gene encoding GPDH enzyme in energy metabolism in *C. pomonella* moth.** (a) RT-qPCR analysis of the expression level of *GPD2* in thorax (a) and leg (b) of male *C. pomonella* moths irradiated with or without 200 Gy X-ray. The data were mean ±SD. The asterisk indicates significant differences as determined by student’s *t*-test (***P*<0.01, ****P*<0.001). n.s. indicates no significant difference.

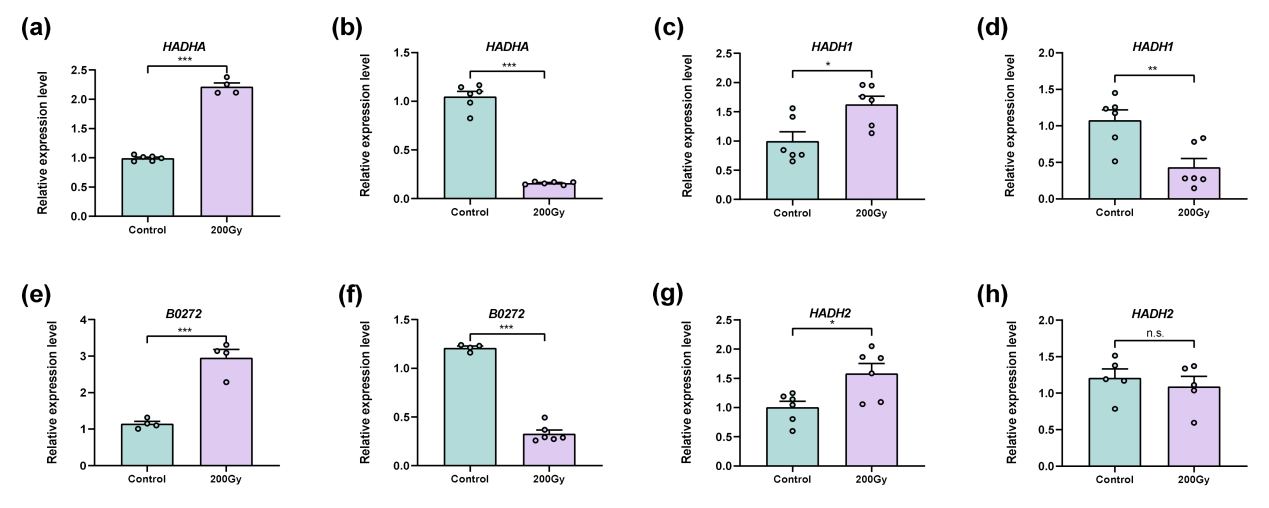

**Figure S4. Effect of 200 Gy X-ray irradiation on expression of genes encoding HOAD enzyme in energy metabolism in *C. pomonella* moth.** RT-qPCR analysis of the expression level of *HADHA* in thorax (a) and leg (b) of male *C. pomonella* moths irradiated with or without 200 Gy X-ray. RT-qPCR analysis of the expression level of *HADH1* in thorax (c) and leg (d) of male *C. pomonella* moths irradiated with or without 200 Gy X-ray. RT-qPCR analysis of the expression level of *B0272* in thorax (e) and leg (f) of male *C. pomonella* moths irradiated with or without 200 Gy X-ray. RT-qPCR analysis of the expression level of *HADH2* in thorax (g) and leg (h) of male *C. pomonella* moths irradiated with or without 200 Gy X-ray. The data were mean ±SD. The asterisk indicates significant differences as determined by student’s *t*-test (**P*<0.05, ***P*<0.01, ****P*<0.001). n.s. indicates no significant difference.
